# Effects of confinement on body mass and site fidelity of feral pigeons during the setting-up of pigeon houses

**DOI:** 10.1101/194043

**Authors:** Julien Gasparini, Lise Dauphin, Justine Favrelière, Adrien Frantz, Lisa Jacquin, Charlotte Récapet, Anne-Caroline Prévot

## Abstract

Feral pigeons can reach high densities in the urban environments and have thus been subject to various regulation programs. Recently, an alternative ethical regulation strategy based on the installation of artificial breeding facilities has been tested in European cities. In Paris (France), pigeons are first confined for several weeks within the pigeon house before being released. According to authorities, this method allows to retain confined pigeons in this new habitat and to attract more conspecifics. This study aims at evaluating the efficiency and potential side-effects of this method by assessing pigeon fidelity behaviour and pigeon welfare after release. Results show that confinement in pigeon houses induced a significant body mass loss in birds. Only 19% of confined pigeons became faithful to their new habitat. This fidelity depended on the origin of birds suggesting that pigeons captured closer to the pigeon houses are more likely to stay in the vicinity of the pigeon house one year after. Investigations on methods of regulation on animal behavior may help to improve management procedures.

---

Urban animals have always coexisted with humans in cities, sometimes for historical reasons, depending on the predominant culture (Clark 2013). Until the beginning of the 20^th^ century, domestic species were dominant in the urban landscape, such as pack animals (horses, cows), livestock (cows, goats, pigs, pigeons) or pets (cats, dogs) (Sabloff 2001). Nowadays, the only domestic species present in the cities are pets, but wild animal species are still present and managed in towns, in the positive vision of urban nature (Matsuoka and Kaplan 2008).

Feral pigeons *Columba livia* have an intermediary status, because they have domestic ancestors but are now thriving in cities independently of human care (Jerolmack 2008; Skandrani et al., 2014). They are present in cities worldwide, and their populations can reach high densities. Feral pigeons and their management procedures may cause social conflicts among citizens because different social groups of citizens often have very strong positioning for or against them (Jerolmack 2008; Colon and Lequarré 2013; Skandrani et al., 2014).Populations of urban pigeons have been subject to public regulation programs in many cities, including culling procedures (reviewed in Haag-Wackernagel 2002). In the last decades, an alternative regulation strategy based on the installation of artificial breeding facilities such as pigeon houses has been tested in European cities to limit local nuisances (e.g. Basel, Switzerland-Haag-Wackernagel 1995; Paris, France-Contassot 2007). Pigeon houses are artificial breeding places where eggs are removed or sterilized using various techniques to limit reproduction with variable consequences for pigeon reproduction and health (Jacquin et al., 2010; Gasparini et al., 2011). Pigeon houses are presented by local authorities as a mean to limit hatching rate and to maintain a small but healthy population (Contassot 2007). This regulation method is also promoted by associations for animal protection, as an ethical method that does not injure or kill individuals (Lizet and Milliet 2012). The management procedures carried out in pigeon houses are variable, but their relative efficiency and side-effects are still unclear, so that few information is available for managers to choose regulation methods in pigeon houses.

In Paris (France), the setting-up of pigeon houses consists in a confinement of a part of the local population during three to four weeks and providing pigeons with food and water.According to authorities and pigeon house managers (Mairie de Paris, 2007; SREP, http://www.srep.fr/), this method is presented as allowing to retain confined pigeons in this new habitat and to attract more conspecifics. However, this confinement may have consequences for pigeons by increasing stress and impacting their health (Wingfield and Romero, 2001). This confinement method is thus perceived as harmful for captive pigeons and is rejected by protection associations. In this study, in agreement with Paris municipality, using an observational approach, we examined the effect of confinement on pigeon fidelity behaviour to the pigeon house and the change in body mass before and after the 3 or 4 weeks of confinements. Body mass loss is a good proxy indicating welfare of individuals (Hawkins 2001; Jacquin et al. 2012). To our knowledge, this is the first scientific study testing the effect of this procedure on pigeon welfare and behaviours.

## Material and methods

The regular implementation of public pigeon houses in Paris belongs to the official program of the current Paris authorities (Mairie de Paris 2007). These public structures are managed by a private company, the Society for Regulation in Pigeon Houses (SREP) which is in charge of attracting pigeons in the structure, of feeding them and controlling reproduction regularly. In agreement with Paris municipality, we performed the study in 2010 and 2011, when three new pigeon houses were implemented: Saint-Eloi (District 12), Saint-Cloud (District 16) and Javel (District 15) in Paris. 32 pigeons were captured at Saint-Eloi, 47 pigeons at Saint-Cloud on the 17^th^ March 2010 and 29 pigeons were captured at Javel on the 13^th^ April 2011 using bait cages with corn. The pigeons in Saint-Eloi and Javel were captured in the vicinity of pigeon houses (in a radius of 50 meters around) while pigeons in Saint-Cloud were captured two kilometers away from the pigeon house with the same baiting cages. After the capture, birds were individually marked with a combination of three color rings that allowed us to identify them from a distance. Pigeons were also weighed to the nearest 5 g with a spring balance (Medio-Line 40600; Pesola, Baar, Switzerland) and an age class was visually determined (juveniles vs. adults) based on the eye color and on the feather shape (Johnson & Janiga 1995). The light sexual dimorphism in pigeons did not enable us to visually sex them, we assumed that sex-ratio did not differ between pigeon houses. In total, 108 pigeons were captured and confined including 10 juveniles and 98 adults. The number of juveniles was significantly more important (Fisher exact test, P = 0.001) in Saint-Eloi (8 over 32) than in Javel (2 over 47) and Saint-Cloud (0 over 29). Pigeons were then confined in the pigeon houses until the 6^th^ April 2010 for Saint-Eloi (i.e., 21 days), until 15^th^ April 2010 for Saint-Cloud (i.e., 30 days) and until 12^th^ May 2011 for Javel (i.e., 29 days). This protocol was constrained by the SREP company. During the confinement, food was supposed to have been provided *ad libitum* by the SREP, with a mix of corn, wheat, and peas supplemented with minerals. Pigeons were weighed again at the end of confinement before being released. For commercial reasons, we were not authorized to follow the protocol performed by the SREP during the confinement period. So we were not able to know whether food and water was effectively provided *ad libitum*. The year following the confinement (between January and March 2011 and 2012, respectively, for Saint-Eloi and Saint-Cloud and for Javel), we looked for marked pigeons once every two weeks during two months (5 sessions of monitoring) around the pigeon houses to monitor their fidelity behaviour. Each of the 5 monitoring session lasted 30 minutes and was performed by two of us (LD and CR). We consider an individual faithful to the pigeon house when it was seen, at least one time during the 5 sessions, within a radius of 20 meters around the pigeon house one year later. It includes birds seen either on the top, on the feet or in front of the exit of the pigeon house. This fidelity, therefore, includes birds that used pigeon house either to eat, to nest or living in its close proximity. This study was carried out in strict accordance with the recommendations of the European Convention for the Protection of Vertebrate Animals used for Experimental and Other Scientific Purposes (revised Appendix A) and with the Guide for the Care and Use of Laboratory Animals. All experiments and captures were approved by local authorities and the “Direction Départementale des Services Vétérinaires de Seine-et-Marne” (permit No. 77-05).

All statistical analyses were then performed on the data using SAS (version 9.4).

## Results

In all three pigeon houses, the number of released pigeons was lower than the number of confined pigeons (Table 1). Most of them were missing and three of them were found dead in the pigeon house at Saint-Eloi. According to the SREP, missing birds escaped during the feeding.

**Table 1:** Numbers and percentages of individuals confined in pigeon houses, released 3 weeks later and seen the following year in pigeon houses.

|  | **Total** | **Saint-Cloud** | **Saint-Eloi** | **Javel** |
| --- | --- | --- | --- | --- |
| **Confined** | 108 (100%) | 47 (100%) | 32 (100%) | 29 (100%) |
| **Released** | 75 (69.4%) | 36 (76.6%) | 21 (65.6%) | 18 (62.1%) |
| **Seen alive one**<br>**year later**<br>**around the**<br>**pigeon house** | 21 (19.4%) | 4 (8.5%) | 9 (34.4%) | 8 (27.6%) |

During the confinement, pigeons lost a significant amount of body mass in the three different pigeon houses (paired student t-test; Saint-Cloud: 7% of body mass loss, t_35_ =4.53, *P*≤ 0.0001; Saint-Eloi: 19% of body mass loss, t_20_ = 10.14, *P* ≤ 0.0001; Javel: 9% of body mass loss, t_17_ = 4.64, *P* = 0.0002; Figure 1). The mass loss differed significantly among pigeon houses (ANOVA, F_2,72_ = 11.36, *P* ≤ 0.0001, Figure 1). Pots-hoc tests revealed that the loss was significantly more important in Saint-Eloi than in Saint-Cloud (Tukey-Kramer, *P*≤0.0001) and in Javel (*P* = 0.0025). The mass lost did not differ between Saint-Cloud and Javel (*P* = 0.85). We compared this body mass loss with body mass changes observed for 112 pigeons captured in the urban environment and placed in captivity in outdoor aviaries in similar food conditions in 2009 (*ad libitum* mix of corn, wheat, and peas supplemented with minerals; Jacquin et al., 2012). Results show that pigeons fed *ad libitum* in captivity gained a significant amount of body mass (15% gain on average) over 30 days (t-test: t_111_ = 7.57, P ≤0.0001).

**Figure 1:**
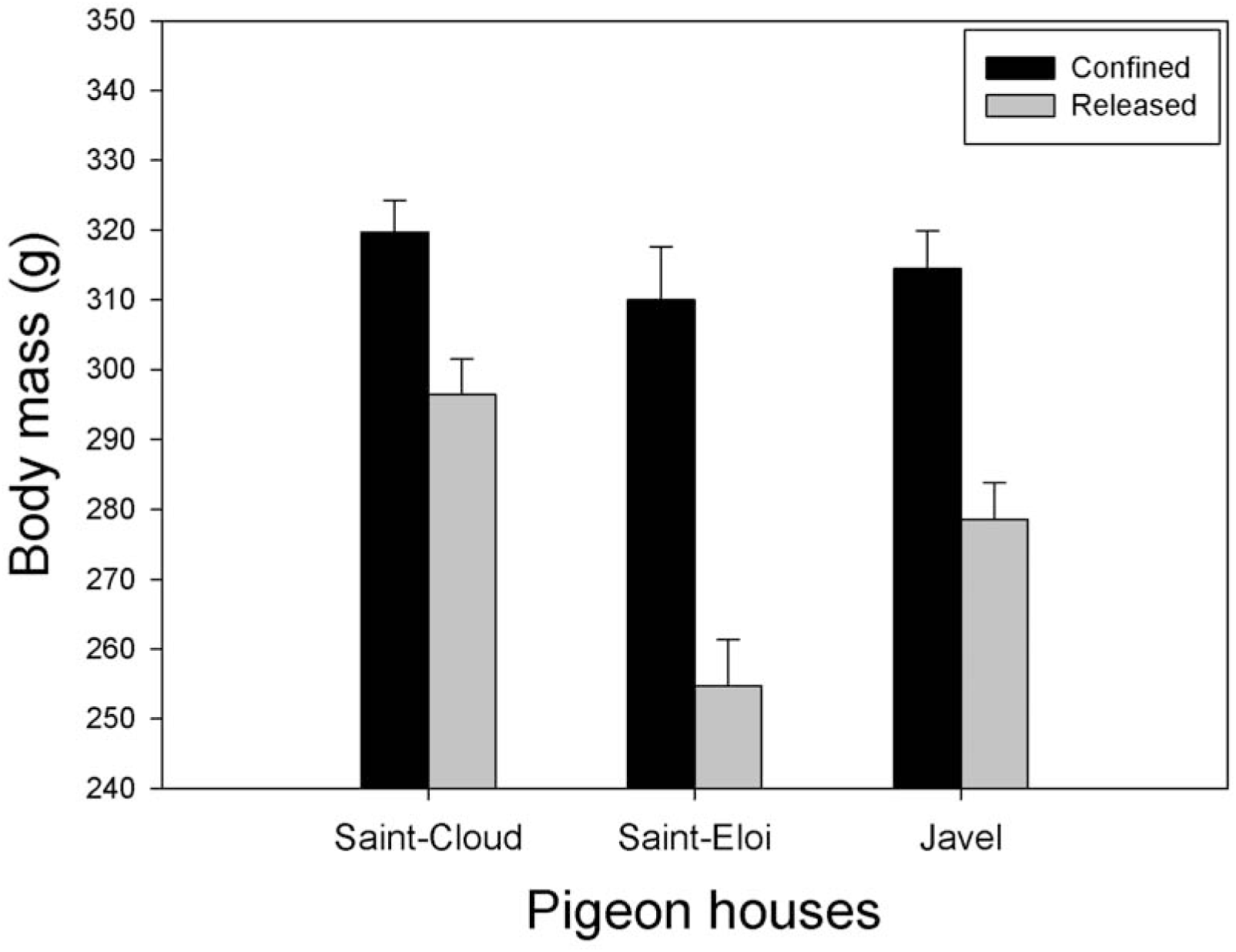
Average body mass ± se of pigeons at time of confinement (black) and at time of releasing (grey) of the three pigeon houses.

One year after the confinement, 19.4 % of pigeons were seen alive and present close to the pigeon houses where they have been confined (Table 1). Ten of these 21 pigeons were seen only once over the 5 sessions, 5 were seen two times, 1 was seen three times and 5 were seen four times. This distribution did not vary significantly among pigeon houses (Fisher exact test *P* = 0.16). However, the proportions of pigeons seen at least one times one year after the confinement significantly differed among pigeon houses (Logistic regression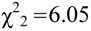, Table 1) but not between ages of pigeon (Logistic regression, 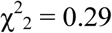, *P* = 0.59). Pots-hoc tests revealed that this proportion was significantly lower in Saint-Cloud than in Saint-Eloi 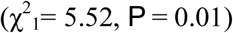 and in Javel 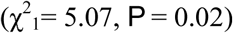. These proportion did not differ between Saint-Eloi and Javel 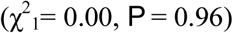. Among the 21 pigeons seen alive one year after the confinement, 18 were adults and 3 juveniles. All re-observed juveniles were from the Saint-Eloi pigeon house. The re-observation rate of juveniles (30.0 %) did not differ from the adult one (18.4 %; Fisher exact test P = 0.40).

## Discussion

This study showed consistent results in three different pigeon houses. In all of them, the confinement of birds within the pigeon houses for 3 weeks strongly reduced their body mass (Figure 1), which could have detrimental effects on their health status (Møller 1998). This loss of body mass could have been caused by captivity. However, another subset of pigeons caught in Paris and put in captivity in open aviary and fed *ad libitum* had a significant increase in their body mass. Several alternative causes can be proposed to explain this strong decrease in body mass after confinement. First, body mass loss may have just resulted from a natural and biological variation in this species (Sargisson et al. 2007). Indeed, there are several examples of seasonal body mass loss such as during the migration, during the incubation or during the chick rearing period (Bryant 1988; Schwilch et al., 2002). However, pigeons are non-migrating birds and no reproduction event occurred during the confinement in our study. Our results are thus not consistent with a natural variation of body mass. Another potential cause of this body mass reduction is the stress caused by confinement conditions. This stress could have been caused by capture and manipulations of the birds before the confinement. Indeed, birds were weighed and marked in the three pigeon houses; moreover, in Saint-Eloi and Saint-Cloud, we took a blood sample for epidemiological analyses (Gasparini et al., 2011), and this manipulation could have been stressful for pigeons and caused the observed decrease of body mass. This interpretation is however unlikely for two reasons: first, a body mass loss was also observed in pigeons in Javel for which no blood sample were taken. Secondly, in another subset of pigeons (Jacquin et al., 2012), captured and manipulated in the urban environment in the same manner, and then put in captivity individually in aviaries with *ad libitum food*, increased their body mass after 30 days (see also Poling et al., 1990).

The last interpretation is that confinement *per se* might dramatically increase the stress for pigeons resulting in a significant decrease in body mass. First, living in a dense group with a unique source of food and increased proximity between individuals could be a factor of elevated stress, as shown in mice (Bartolomucci et al., 2004). Pigeon is known to be a social species with a strong hierarchical structure (Johnston and Janiga, 1995; Sol et al., 1998). The fact that food was provided in a unique localization may increase competition within the pigeon house and may increase aggressive interactions among individuals. Second, the confinement may induce a psychological stress responsible of body mass loss (Morgan and Tromborg, 2007). As we were not able to check that food was providing *ad libitum* during the confinement, we also cannot fully exclude that food was not lacking during this period Finding the mechanisms responsible of this body mass loss would allow us to find alternative methods to avoid negative side-effects of pigeon houses on pigeon condition.

A second interesting result of our study is the estimation of fidelity of confined birds. The confinement enabled for approximately 19% of birds to become faithful to this new habitat. This low fidelity is however difficult to interpret for several reasons. First, when a bird is not re-observed one year after, it might be dead or have migrated to another site. Therefore, with our method, we cannot distinguish between mortality and fidelity. In any cases, the objective of the pigeon houses is to fix alive individuals in these latter and, according to our results, this objective is only few filled (only for 19% birds) either because the confinement induced high mortality or did not enable to make birds loyal enough to the pigeon house. Alternatively, the low site fidelity observed in our study might be caused by the egg removing that reduce reproductive success. Indeed, previous studies on habitat selection predict that reproductive success may directly impact whether individuals are coming back to the same site of reproduction or are leaving to another one (Switzer 1993). Future studies should therefore investigate how egg removal may alter the site fidelity. Interestingly, the re-observation rate in the pigeon houses of Saint-Eloi and Javel were higher (around 30 %) than the re-observation in the Saint-Cloud pigeon house (8.5 %, Table 1). Contrary to Saint-Eloi and Javel, the pigeons confined in Saint-Cloud were not captured on the site of the pigeon house, but 2 km away from it. Although further studies need to be carried out to confirm our results, this suggests that pigeons should be captured very close to the site of the pigeon house to ensure a long-term fidelity to the pigeon house. Indeed, several studies outlined the limited home range of pigeons in the urban environment (mostly below 1.5 km of radius; Sol and Senar 1995; Rose et al., 2006; Frantz et al., 2012), so that capturing pigeons close to the pigeon house could prevent their return to their previous home range and increase the chance of the setting-up of a permanent and healthy pigeon colony within the pigeon house. Alternatively, the higher re-observation rate in Saint-Eloi could be due to the higher frequency of juveniles confined in this pigeon house. However, this effect was not significant and, therefore, this interpretation is unlikely.

## Conclusion

In conclusion, our study is the first to evaluate the method of setting-up of pigeon houses for regularization purposes. Our results showed that the confinement before the opening of the pigeon house has dramatic consequences for birds in terms of body mass loss with potential negative consequences for their health status and survival, although the underlying mechanism remains to be identified. Results also suggest that the fidelity of confined birds to the pigeon house after one year depends on the origin of birds and might be improved by local captures around the pigeon house.

The implementation of pigeon houses to manage pigeon populations is fully acceptable for pigeons’ protection associations (Lizet and Milliet, 2012), contrary to other public measures such as feeding ban (Colon and Lequarré, 2013). If adequately managed, pigeon houses could therefore serve as mediators between conflicting social groups of the “pigeon problem” (see actor-network theory, Latour 2005). However, to let them play such a role, a high level of quality and transparency in all management decisions is needed, in order to encourage communication and participation in management decision-making. We therefore encourage the communication of the scientific data provided by this study to all managers and stakeholders, to help designing and improving co-management procedures of pigeon populations and ensure a peaceful cohabitation of nature and citizens within the urban habitat.

## Acknowledgements

We thank Paris municipality to have allowed us to survey pigeons in the three studied pigeon houses. We are very grateful to Gérard Leboucher, Hélène Corbel and Philippe Lenouvel for the help they provided at different stages of the study. This work was financed by grants from the local government (Ile-de-France: Sustainable Development Network R2DS, No. 2008-07).Jacquin was supported by a Ph.D. fellowship from the French Ministry of Research. L. Dauphin was supported by the ATM “Relations hommes-nature sur le long terme”, from the national museum of natural history.

## References

Bartolomucci, A., Pederzani, T., Sacerdote, P., Panerai, A.E., Parmigiani, S. and Palanza, P. (2004) Behavioral and physiological characterization of male mice under chronic psychosocial stress. Psychoneuroendocrino., 29, 899–910.

Bryant, D.M. (1988) Energy expenditure and body mass changes as measures of reproductive costs in birds. Funct. Ecol., 2, 23–34.

Clark, P. (2013) Handbook of cities in world history. Oxford University Press, Oxford.

Colon, P.L. and Lequarré, N. (2013) Le nourrissage des pigeons dans la région parisienne. Ethnologie française, XLIII, 153-[In French.]

Contassot, Y. (2007) La politique de la ville : pour une gestion durable des pigeons à Paris, in : Mairie de Paris (Ed.), Bien vivre avec les animaux à Paris, le guide de l'animal en ville. Mairie de Paris, Paris, pp. [In French.]

Frantz, A., Pottier, M.A., Karimi, B., Corbel, H., Aubry, E., Haussy, C., Gasparini, J. and Castrec-Rouelle, M. (2012) Contrasting levels of heavy metals in the feathers of urban pigeons from close habitats suggest limited movements at a restricted scale. Environ. Pollut., 168, 23–28.

Gasparini, J., Erin, N., Bertin, C., Jacquin, L., Vorimore, F., Frantz, A., Lenouvel, P. and Laroucau, K. (2011) Impact of urban environment and host phenotype on the epidemiology of Chlamydiaceae in feral pigeons. Environ. Microbiol., 13, 3186–3193.

Haag-Wackernagel, D. (1995) Regulation of the street pigeon in Basel. Wildlife Society Bulletin, 23, 256–260.

Haag-Wackernagel D (2002) Feral Pigeons: Management Experiences in Europe. In: Dinetti, M.Ed., Atti 2° Convegno Nazionale sulla Fauna Urbana, Specie ornitiche problematiche:Biologia e gestione nelle cittá e nel territorio. pp. 25-37, 10 Giugnio 2000, ARSIA e LIPU., Firenze (Toscana).

Hawkins, P. (2001) Laboratory birds, refinements in husbandry and procedures. Fifth report of the BVAAWF/FRAME/RSPCA/UFAW Joint Working Group on Refinement. Lab. Anim., 35,1–163.

Jacquin, L., Cazelles, B., Prévot-Julliard, A-C., Leboucher, G. and Gasparini, J. (2010) Reproduction management in pigeon houses affects breeding ecology of feral pigeons: implications for control policies. Can. J. Zool., 88, 781–787.

Jacquin, L., Recapet, C., Bouche, P., Leboucher, G. and Gasparini, J. (2012) Melanin-based coloration reflects alternative strategies to cope with food limitation in pigeons. Behav. Ecol., 23, 907–915.

Jacquin, L., Blottiere, L., Haussy, C., Perret, S. and Gasparini, J. (2012) Prenatal and postnatal parental effects on immunity and growth in ‘lactating’ pigeons. Funct. Ecol., 26, 866– 875.

Jerolmack, C. (2008) How pigeons became rats: the cultural-spatial logic of problem animals.Social Problems, 55, 72–94.

Johnston, R.F. and Janiga, M. (1995) Feral pigeons. Oxford University Press, Oxford.

Latour, B. (2005) Reassembling the Social: An Introduction to Actor-Network-Theory. Oxford University Press, Oxford.

Lizet, B. and Milliet, J. (2012) Le pigeonnier public, à la croisée des utopies sur le vivant dans la ville. In : Lizet, B. and Milliet J. eds, « Animal certifié conforme. Déchiffrer nos relations avec le vivant », pp. 185–204. Dunod, Paris. [In French.]

Mairie de Paris. Bien vivre avec les animaux à Paris, protégeons notre environnement. Dossier Environnement. http://www.paris.fr/publications/publications-de-la-ville/guides-et-brochures/bien-vivre-avec-les-animaux-a-paris/rub_6403_stand_15553_port_14438. [In French.] [accessed 16 January 2016].

Matsuoka, R.H. and Kaplan, R. (2008) People needs in the urban landscape: analysis of Landscape and Urban Planning contributions. Landscape Urban Planning, 84, 7–19.

Møller, A.P., Christe, P., Erritzøe, J. and Mavarez, J. (1998) Condition, disease and immune defence. Oïkos, 83, 301–306.

Morgan, K.N. and Tromborg, C.T. (2007) Sources of stress in captivity. Appl. Anim. Behav. Sci., 102, 262–302.

Poling, A., Nickel, M. and Alling, K. (1990) Free birds aren't fat: weight gain in captured wild pigeons maintained under laboratory conditions. J. Exp. Anal. Behav., 53, 423–424.

Rose, E., Nagel, P. and Haag-Wackernagel, D. (2006) Spatio-temporal use of the urban habitat by feral pigeons (*Columba livia*). Behav. Ecol. Sociobiol., 60, 242–254.

Sabloff, A. (2001) Reordering the natural world. Humans and animals in the city. University of Toronto Press, Toronto, Canada.

Schwilch, R., Grattarola, A., Spina, F. and Jenni, L. (2002) Protein loss during long distance migratory flights in passerine birds: adaptation and constraint. J. Exp. Biol., 205, 687– 695.

Sargisson, R.J., McLean, I.G., Brown, G.S. and White, K.G (2007) Seasonal variation in pigeon body weight and delayed matching-to-sample performance. J. Exp. Anal. Behav., 88: 395–404.

Skandrani, Z., Lepetz, S. and Prévot-Julliard, A-C. (2014) Nuisance species: beyond the ecological perspective. Ecological Processes, 3, 3.

Sol, D., Santos, D.M., Garcia, J. and Cuadrado, M. (1998) Competition for food in urban pigeons: the cost of being juvenile. Condor, 100, 298–304.

Sol, D. and Senar, J.C. (1995) Urban pigeon populations: stability, home range, and the effect of removing individuals. Can. J. Zool., 73, 1154–1160.

Switzer, P. V. (1993) Site fidelity in predictable and unpredictable habitats. Evol. Ecol., 7, 533–555.

Wingfield, J.C. and, Romero, L.M. (2001) Adrenocortical responses to stress and their modulation in free-living vertebrates. In: McEwen, B.S. ed. Handbook of physiology. pp. 211–234, Oxford University Press, New York.

